# Regulatory role of cholesterol in modulating actin dynamics and cell adhesive interactions in the trabecular meshwork

**DOI:** 10.1101/2024.02.02.578717

**Authors:** Ting Wang, Hannah R C Kimmel, Charles Park, Hyeon Ryoo, Jing Liu, Gregory H Underhill, Padmanabhan P Pattabiraman

**Affiliations:** Glick Eye Institute, Department of Ophthalmology, Indiana University School of Medicine, 1160 West Michigan Street, Indianapolis, Indiana, 46202, United States of America; Stark Neuroscience Research Institute, Medical Neuroscience Graduate Program, Indiana University School of Medicine, 320 W. 15th Street, Indiana, 46202, United States of America; Department of Bioengineering, University of Illinois at Urbana-Champaign, Urbana, Illinois, 61801, United States of America; Deparment of Physics and Astronomy, Purdue University, 525 Northwestern Avenue, West Lafayette, Indiana, 47907, United States of America

**Keywords:** Cholesterol, Trabecular meshwork cell, Actin cytoskeleton, Integrin, Phosphatidylinositol 4, 5-bisphosphate

## Abstract

The trabecular meshwork (TM) tissue plays a crucial role in maintaining intraocular pressure (IOP) homeostasis. Increased TM contractility and stiffness are directly correlated with elevated IOP. Although cholesterol is known to be a determinant of glaucoma occurrence and elevated IOP, the underlying mechanisms remain elusive. In this study, we used human TM (HTM) cells to unravel the effects of cholesterol on TM stiffness. We achieved this by performing acute cholesterol depletion with Methyl-β-cyclodextrin (MβCD) and cholesterol enrichment/replenishment with MβCD cholesterol complex (CHOL). Interestingly, cholesterol depletion triggered notable actin depolymerization and decreased focal adhesion formation, while enrichment/replenishment promoted actin polymerization, requiring the presence of actin monomers. Using a specific reporter of phosphatidylinositol 4,5-bisphosphate (PIP2), we demonstrated that cholesterol depletion decreases PIP2 levels on the cell membrane, whereas enrichment increases them. Given the critical role of PIP2 in actin remodeling and focal adhesion formation, we postulate that cholesterol regulates actin dynamics by modulating PIP2 levels on the membrane. Furthermore, we showed that cholesterol levels regulate integrin α5β1 and αVβ3 distribution and activation, subsequently altering cell-extracellular matrix (ECM) interactions. Notably, the depletion of cholesterol, as a major lipid constituent of the cell membrane, led to a decrease in HTM cell membrane tension, which was reversed upon cholesterol replenishment. Overall, our systematic exploration of cholesterol modulation on TM stiffness highlights the critical importance of maintaining appropriate membrane and cellular cholesterol levels for achieving IOP homeostasis.

## 1. Introduction

Glaucoma is an age-related optic neuropathy and is one of the leading causes of irreversible blindness (1). Primary open-angle glaucoma (POAG) is the most predominant subtype. Elevated intraocular pressure (IOP) is a major risk factor of POAG, and chronic elevation in IOP damages the optic nerve leading to irreversible blindness (1, 2). Previous studies have found that cholesterol and its biogenic pathway are closely related to POAG occurrence and IOP elevation. Glaucoma patients showed a significantly higher total cholesterol levels than patients without glaucoma (3–5) and meta-analysis reported that increased cholesterol levels are associated with a higher risk of glaucoma and IOP elevation (6). Our recent work provided evidence that the inactivation of the transcription factors sterol regulatory element binding proteins (SREBPs), involved in fatty acid and cholesterol biogenesis, significantly lowered intraocular pressure (7). Moreover, studies found that the use of statins, cholesterol biosynthesis inhibitors, can reduce the incidence and progression of glaucoma (8–12) in spite of some inconsistencies regarding the role of cholesterol in POAG pathogenesis (13). Although cholesterol is related to glaucoma pathogenesis and IOP elevation, the causal effects of cholesterol in IOP elevation and glaucoma progression are not fully understood.

Currently, lowering IOP is the most effective way to delay and halt the progression of POAG (1). The IOP within the eye is maintained by the balance of aqueous humor (AH) generated by the ciliary body and drainage via the conventional outflow pathway consisting of trabecular meshwork (TM), juxtacanalicular TM tissue (JCT), and Schlemm’s canal (SC) (14). Strong experimental evidence demonstrates that increased actin contractility and ECM-mediated stiffness in TM are directly related to IOP elevation and POAG pathogenesis (15–19). Additionally, cell-ECM interactions via integrins are involved in regulating TM biomechanics and IOP (20–23). Cholesterol is the major sterol component of the cell membrane, which makes up about 30-40% of the lipid bilayer on average (24). It is an important player in modulating various cellular signaling pathways, membrane properties, actin dynamics, and cell-ECM interactions (25, 26). Cholesterol levels directly regulate lipid order in the cell membrane (27), as well as membrane tension and fluidity (28). Increased cholesterol content on the cell membrane significantly increases the order of the lipid packing (29), lowers the membrane permeability (30, 31), decreases membrane fluidity, and increases rigidity (32). Cholesterol is the principal component of membrane microdomains such as caveolae and lipid rafts serving as important locations to anchor transmembrane proteins, actin cytoskeleton binding, and cell-ECM communications (33). Interactions between microdomains and cytoskeletal components can contribute to regulating microdomain assembly as well as cytoskeletal dynamics. Actin-binding proteins and other focal adhesion (FA) proteins are localized to microdomains, thus providing a platform for cytoskeletal tethering and cell-ECM interactions through integrins, cadherins, occludins, and other cellular adhesion molecules (33). As a major component of microdomains, cholesterol levels on the cell membrane have a direct effect on modulating actin-membrane binding and integrins signaling in microdomains (34–36). Although these effects of cholesterol have been studied in multiple cell types and artificial lipid membranes, the detailed mechanisms, and effects of cholesterol on TM contractility and stiffness are still blurred. Earlier studies found that statins can lower the IOP and increase AH outflow facility in animal models by suppressing Rho isoprenylation, Rho-kinase activity, and actin polymerization in the TM (37–39). However, one study found that the removal of cholesterol in human TM (HTM) cells using methyl beta cyclodextrin (MβCD) caused increased actin stress fibers and augmented the activation of the transient receptor potential vanilloid isoform 4 (TRPV4) channels (40). Interestingly though, our recent study demonstrated a decrease in cholesteryl ester levels in HTM cells after mechanical stretch (18). Cholesteryl esters are the storage form of cholesterol, and their decrease may indicate increased free cholesterol levels in HTM cells under mechanical stretch. Further, the activation of SREBP2, which increases cholesterol biosynthesis, can result in increased actin polymerization and ECM accumulation in HTM cells (7). Yet the direct correlation between cholesterol levels, actin-cell adhesive interactions, and TM biomechanics is not available. This study deciphers the mechanistic evidence for the regulation of actin polymerization, actin-membrane binding, cell-ECM interaction, and cell membrane tension and fluidity in TM cells upon perturbing cellular and membrane cholesterol levels.

## 2. Material and Methods

### 2.1 Materials

Materials are listed in **Table 1**.

**Table 1.**
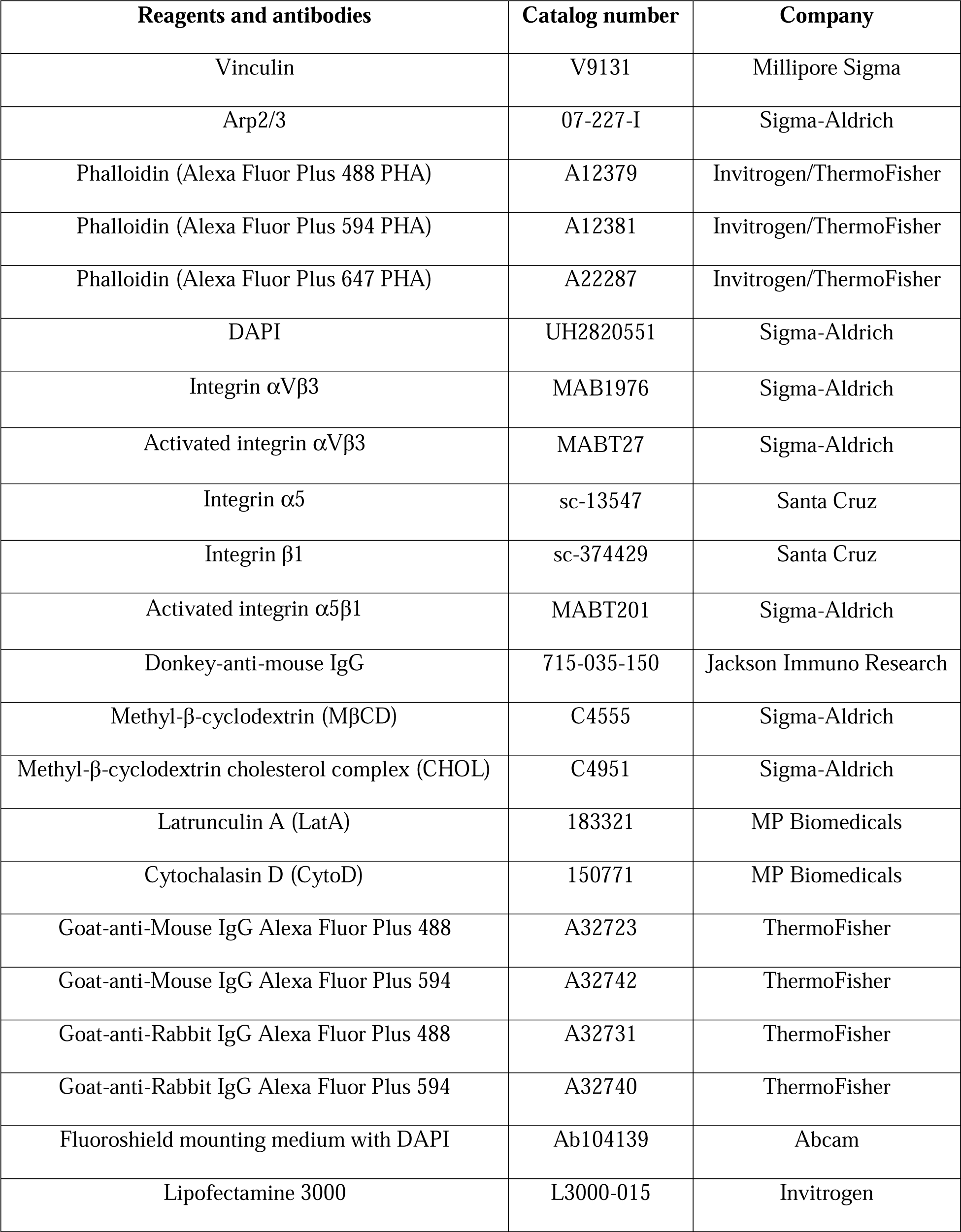
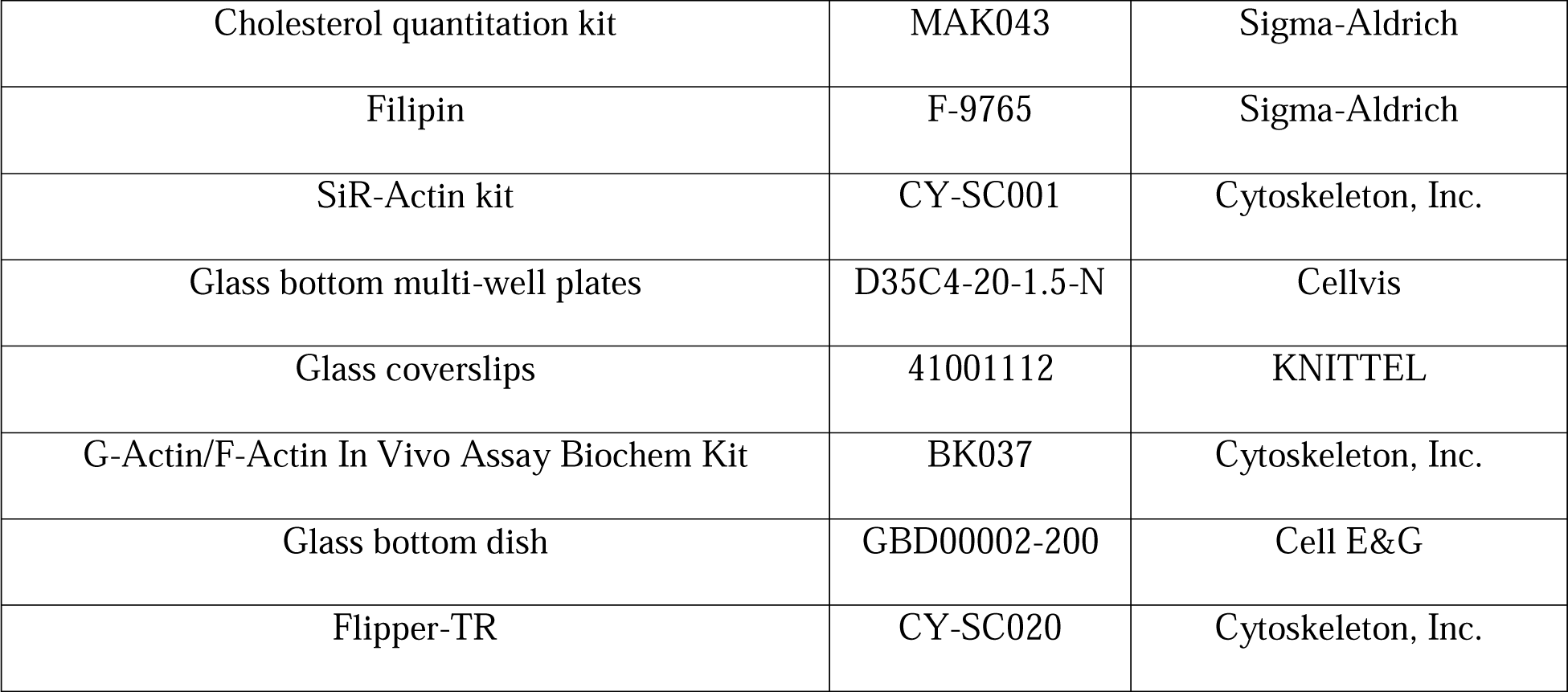
Materials.

### 2.2 TM cell cultures

Primary HTM cells were cultured from TM tissue isolated from the leftover donor corneal rims after they had been used for corneal transplantation at the Indiana University Clinical Service, Indianapolis. HIPPA compliance guidelines were adhered to for the use of human tissues. The usage of donor tissues was exempt from the DHHS regulation and the IRB protocol (1911117637), approved by the Indiana University School of Medicine IRB review board. Four lines of the HTM cells used in this study were characterized using dexamethasone induced-myocilin expression (41). Immortalized normal human TM (NTM) cell lines were generously provided by Abbot Clark, University of North Texas Eye Research Institute, Dallas, Texas. All experiments were conducted using confluent or semi confluent TM cultures as mentioned. HTM cells were used between passages four to six. Experiments were performed after overnight serum starvation unless mentioned otherwise. All experiments utilizing HTM cells in this manuscript were performed using biological replicates.

### 2.3 Cell culture treatments

Overnight serum starvation was carried out on cell cultures before treatments. Cholesterol depletion was achieved by using 10 mM MβCD treatment for 1 h. Cholesterol enrichment was achieved by using 100 μM CHOL treatment for 1 h. Cholesterol replenishment was done after 10 mM MβCD treatment for 1 h and cells were washed with serum-free media three times to completely remove MβCD, then were treated with 100 μM CHOL for 1 h to replenish cholesterol (**Figure 1A**). Cholesterol biosynthesis inhibition was done using 100 μM atorvastatin treatment for 24 h. Inhibition of actin polymerization was achieved by treating HTM cells with 10 μM cytochalasin D (CytoD) and 2 μM Latrunculin A (LatA) for 1 h. For CytoD and LatA in combination with CHOL treatments, cells were first treated with CytoD or LatA for 1 h followed by treatment with CHOL without washing for another 1 h to check for the effects of cholesterol on actin polymerization.

**Figure 1.**
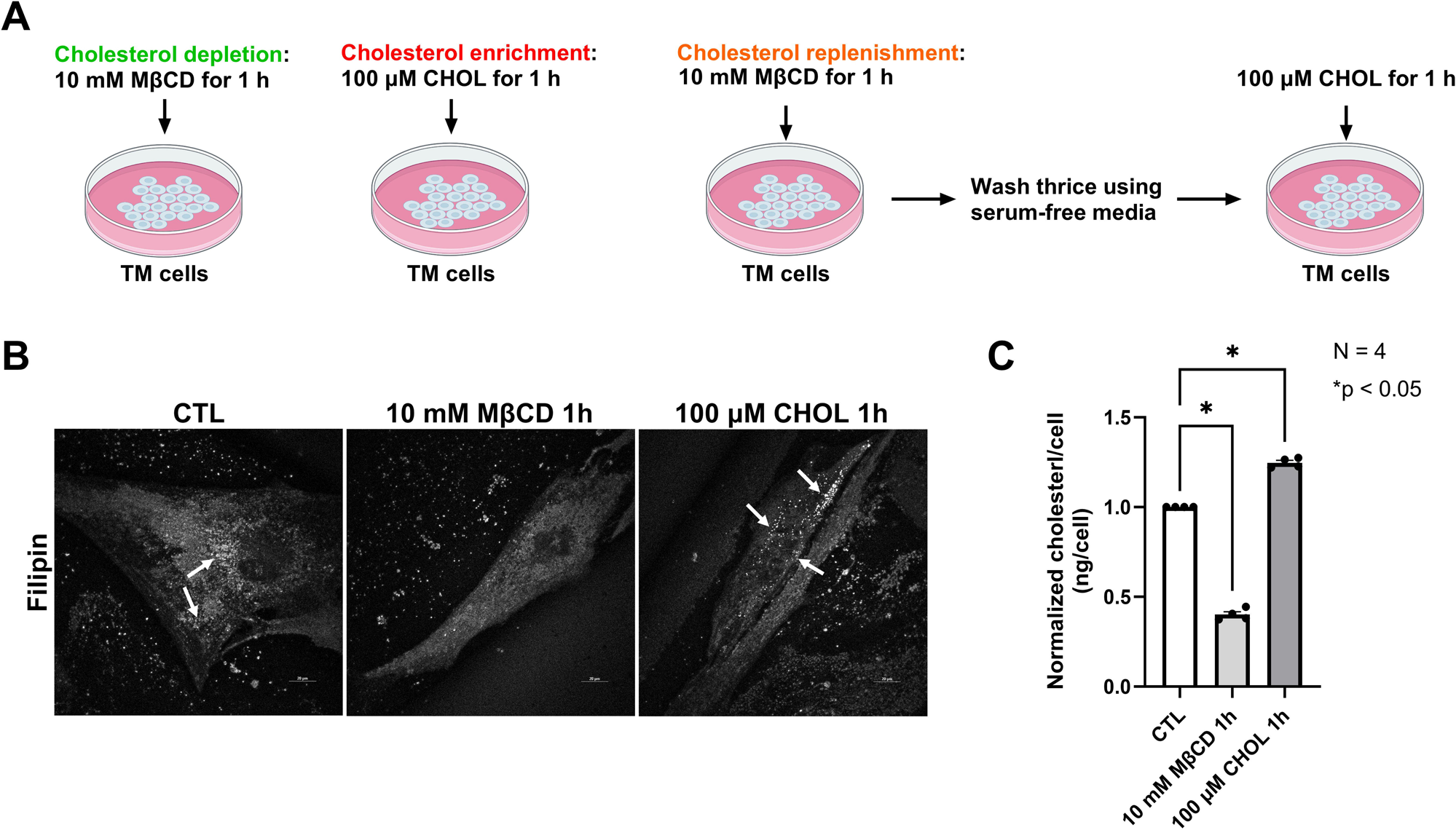
MβCD and CHOL modulate cholesterol levels in HTM cells. (A) The experimental paradigm utilized in the study. (B) Free cholesterol contents in HTM cells were imaged using filipin staining. Under the control condition, cholesterol puncta were identified (denoted by white arrows). Compared to the control condition, cholesterol depletion by 10 mM MβCD treatment for 1 h decreased cholesterol distribution, but cholesterol enrichment by 100 μM CHOL treatment for 1 h increased cholesterol distribution along with more cholesterol puncta observed in HTM cells (denoted by white arrows). (C) Free cholesterol levels in HTM cells were measured using a cholesterol quantification kit. Compared to the control condition, cholesterol depletion by MβCD significantly lowered the cholesterol levels, and cholesterol enrichment by CHOL significantly increased cholesterol levels. Values represent the mean ± SEM, where n = 4 (biological replicates). *p<0.05 was considered statistically significant. CTL: control condition.

### 2.4 pEGFP-N1-Tubby(c)R332H plasmid transfection

Tubby protein contains a phosphatidylinositol 4,5-bisphosphate (PIP2) binding domain (tubby domain). It has been reported that PIP2 depletion aids the release of the full-length tubby protein from the cell membrane and nuclear translocation (42). Because of this selective binding domain, it was reported that tubby is a useful probe to examine changes in PIP2 in intact cells (43, 44). To better understand if cholesterol regulates PIP2 levels in TM cells, NTM cells were transfected with pEGFP-N1-Tubby(c)R332H using Lipofectamine 3000 as per manufacturer protocol. Post 24 h, cells were treated with 10 mM MβCD or 100 μM CHOL for 1 h, stained for actin using SiR-Actin, and imaged using TIRF microscopy. pEGFP-N1-Tubby(c)R332H was a generous gift from Dr. Gerry Hammond, University of Pittsburgh.

### 2.5 Cholesterol content measurement and imaging

HTM cells were plated on 100 mm plates. Post-treatment (10 mM MβCD or 100 μM CHOL for 1 h), cells were counted (around 0.15-0.6 x 10^6^), and were collected and frozen at −80°C until analysis. Then the free cholesterol from HTM cells was extracted and measured with a cholesterol quantitation kit, following the manufacturer’s instructions.

For imaging cholesterol content, HTM cells plated on coverslips were treated with 10 mM MβCD or 100 μM CHOL for 1 h and rinsed thrice with 1x PBS. Later cells were fixed with 4% paraformaldehyde, washed thrice with PBS and incubated with 1 ml of 1.5 mg glycine/ml PBS for 10 min at room temperature to quench the paraformaldehyde. Then the cells were stained with 0.05 mg/ml filipin for 2 h at room temperature. After rinsing cells thrice with PBS, cells were imaged using a Nikon AX/AX R confocal microscope system.

### 2.6 Live cell imaging

Live-cell imaging for observing changes in filamentous actin (F-actin) was performed using a SiR-Actin kit. HTM cells were grown on 2% gelatin-coated glass bottom multi-well plates. When cells reached the desired density, the actin staining solution containing 1 μM SiR-Actin and 10 μM verapamil was added. The cells were maintained in the incubator at 37 °C in a humidified atmosphere containing 5% CO_2_. After 12 h, live-cell imaging was performed at 37 °C in a Tokai Hit Stage Top incubator to record the changes in actin structures under a Zeiss LSM 700 confocal microscope. The z-stack images were obtained and processed using Zeiss ZEN image analysis software. The live cell imaging video was processed using ImageJ software (version 1.53a).

### 2.7 Protein distribution analysis by immunofluorescence (IF)

IF staining was performed on HTM cells grown on 2% gelatin-coated glass coverslips. The methodologies used for IF have been published from the lab earlier (7, 17, 45). All the slides were observed under a Zeiss LSM 700 confocal microscope, and z-stack images were obtained and processed using Zeiss ZEN image analysis software.

### 2.8 Total internal reflection fluorescence (TIRF) microscopy imaging

NTM cells were grown on 2% gelatin-coated glass coverslips. When cells reached the desired density, cells were transfected with pEGFP-N1-Tubby(c)R332H plasmid. Post 48 h, cells were washed with 1X PBS twice, fixed in 4% paraformaldehyde for 20 min, permeabilized with 0.2% triton-X-100 in PBS buffer for 10 min, then washed 3 times with 1X PBS, and incubated with phalloidin for 1 h at room temperature. Finally, the coverslips were washed and mounted onto glass slides with Fluoroshield Mounting Medium. All the slides were observed under a Zeiss Axiovert 200M configured with TIRF microscopy.

### 2.9 Quantitative image analysis

ImageJ (version 1.53a) software was used to analyze the F-actin fluorescence intensity in IF images. Specifically, the immunofluorescence images obtained from the cell culture were converted into an 8-bit image, then the threshold default setup under Adjust was used to convert the images from grayscale into a binary image. Next, the region of interest (ROI) tool was chosen for analysis. In all the analysis, random (different) ROIs (equal area) in the cell cultures were chosen in an image and the intensity was measured and compared in a double blinded manner. Once data was plotted, the control and treatment were revealed at the end of the study. Similarly, each F-actin fiber intensity was chosen using the ROI tool in an image, and the intensity was measured and compared between the control and treatment. Similarly, PIP2 fluorescent intensity in the cell membrane in TIRF images was also measured using the ROI tool and compared between the control and treatment.

### 2.10 Quantitative filamentous actin/globular actin (F-actin/G-actin) ratio measurement

The F-actin/G-actin ratio was measured using the G-Actin/F-Actin In Vivo Assay Biochem Kit, following the manufacturer’s instructions and our previously published work (7).

### 2.11 High throughput traction force microscopy (TFM), imaging processing, and microarray analysis

#### 2.11.A Preparation of polyacrylamide hydrogels

Polyacrylamide gels were prepared based on previously published protocol (46). Briefly, glass bottom dishes (35 mm) were washed with 0.25% v/v Triton X-100 in distilled water (dH_2_O) on an orbital shaker for 30 minutes. After rinsing with dH_2_O, dishes were then submerged in acetone and washed for 30 minutes on the shaker. Next, dishes were submerged in methanol and washed for 30 minutes on the shaker. The glass area of each dish was then etched with 0.2 N NaOH for 1 h, rinsed with dH_2_O, dried with compressed air, and placed on a hot plate at 110 °C until completely dry. To silanize the dishes, the glass area of each dish was submerged in 2% v/v 3-(Trimethoxysilyl) Propyl Methacrylate (3-TPM) in ethanol and placed on the shaker to react for 30 minutes. The silanized dishes were then washed with ethanol on the shaker for 5 minutes, dried with compressed air, and again placed on the hot plate at 110 °C until fully dry. To fabricate polyacrylamide hydrogels with defined elastic moduli, a prepolymer solution for 6 kPa Young’s modulus was prepared (6% acrylamide, 0.45% bis-acrylamide). The prepolymer solution was mixed with a 20% w/v solution of Irgacure 2959 (BASF, Corp.) in methanol to achieve a final volumetric ratio of 9:1 (prepolymer:Irgacure). 100 μl of this working solution was then placed onto the silanized glass area of the dish and sandwiched with a coverslip. 20 μl working solution was used for 12 mm dish gels. The dishes containing working solution and coverslips were then transferred to a UV oven and exposed for 10 minutes to 365 nm UVA light (240 x 10^3^ μJ). After polymerization, the coverslips were removed from the gel and dishes were submerged in dH_2_O at room temperature for 72 hours to remove excess reagents from the hydrogels. Before microarray fabrication, hydrogel substrates were dehydrated on a hot plate for at least 15 minutes at 50 °C.

#### 2.11.B Array fabrication

Cellular ECM microarrays were fabricated to perform TFM using previously published protocols (46, 47). Briefly, ECM proteins – Collagen 1 (COL 1), Collagen IV (COL 4), fibronectin (FN), and laminin for arraying were diluted in a 2X printing buffer solution comprised of 40% v/v glycerol and 0.5% v/v Triton X-100 in dH_2_O, 16.4 mg/ml sodium acetate, 3.72 mg/ml EDTA, and subsequently deposited in a 384-well V-bottom microplate. All single ECM solutions were prepared at a final concentration of 250 μg/ml per ECM, with dH_2_O as the diluent to achieve a final volume of 10 μl per microwell. To transfer ECM condition solutions from the source plate to polyacrylamide hydrogel substrate, a robotic benchtop microarrayer (OmniGrid Micro, Digilab) loaded with SMPC Stealth microarray pins (Arraylt) was used, which produced array islands of ∼600 μm diameter. After fabrication, arrays were stored at room temperature and 65% relative humidity overnight and then left to dry at room temperature and standard humidity in the dark. Before adding cells, all arrays were sterilized with 30 minutes UVA while submerged in a 1X PBS solution containing 1% v/v penicillin/streptomycin.

#### 2.11.C Seeding HTM cells onto microarrays for treatments

Three HTM lines (biological replicates) were seeded onto the microarrays. Cells were collected and counted and then resuspended in culture media at an appropriate concentration for seeding. 3 ml cell suspension was added to each dish, and the dishes were incubated at 37 °C and 5% CO_2_ for 24 h until confluent cell islands had formed. Once islands had formed, the arrays were rinsed twice with 3 ml prewarmed media and then were divided into 3 groups in each cell line: control condition, 10 mM MβCD for 1 h, and 100 μM CHOL for 1 h. After treatments, the dishes were imaged using a Zeiss LSM 700 confocal microscope.

#### 2.11.D Traction force microscopy (TFM) imaging, image processing, and microarray analysis

Cellular microarrays were imaged using Zeiss Axiovert 200M. Printed dextran rhodamine fluorescent markers were used to orient the widefield imaging regions and set the spacing of islands in the microarray. Using phase contrast, the XY positions of the cell islands were located and marked. For each XY location, the Z position was set using the verify positions tool such that the top plane of the hydrogel with the fluorescent beads was just in focus. The saved XY positions of the islands were recalled and modified for subsequent dishes. After setting up, the pre-dissociation phase contrast and fluorescent images were captured for each cell island. Then 150 μl SDS solution was carefully added to the dish to dissociate the cell islands from the substrate completely, and then the automated imaging of the islands was repeated to capture the post-dissociation fluorescent images. This process was repeated for each dish. The images were processed to estimate displacement fields and calculate traction fields using the MATLAB code mentioned in previous publications (46, 47). The phase contrast image was used to determine the island location and dimensions for the estimates and calculations. Each island was designated with the stiffness of the hydrogel and its ECM protein makeup due to its position relative to the dextran rhodamine markers.

### 2.12 Fluorescence lifetime imaging microscopy (FLIM) imaging

HTM cells were grown on 2% gelatin-coated glass bottom multi-well plates. After achieving the desired confluency, HTM cells were treated with 1 μM Flipper-TR diluted from 1 mM master stock solution. Flipper-TR is a live cell fluorescent membrane tension probe that specifically inserts into the cell plasma membrane (48). The cells were then returned to the incubator at 37 °C in a humidified atmosphere containing 5% CO_2_ for 15 minutes before imaging. After incubation, the Flipper-TR fluorescent lifetime before treatment, after cholesterol depletion using MβCD treatment for 1 h, and after cholesterol replenishment using CHOL treatment for another 1 h, were recorded using FLIM.

The fluorescence lifetimes of the Flipper-TR tension sensor were measured using a home-built FLIM system based on a laser confocal microscope (FV1000, Olympus) and time-resolved single photon counting. A pulsed laser at 450 nm wavelength (LDH-D-C-450, Picoquant) was used as the excitation light source, which was coupled to the laser scanning module of the confocal microscope; the fluorescent emission was collected by a photon counting module (PDM series, Micro Photon Devices) coupled with a time-correlated single photon counting (TCSPC) module (TimeHarp 260, Picoquant). The fluorescent photons were recorded in the manner of single photon counting, which resulted in a histogram of the number of the emitted photons in terms of the arrival times. For a specified region of interest (ROI), the histogram was fitted by a two-component exponential decay function (49).

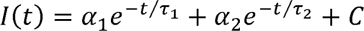

*I* is the *t*-dependent emission intensity of the target fluorophores, whereas τ*_i_* is the *i*-th component of the decay lifetime with the corresponding fraction of α*_i_*. *C* is a constant used to compensate for the background noise of the measured signal. The average fluorescence lifetime of the signal is tabulated using the τ*_i_*’s and α*_i_*’s as in the following:

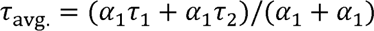

The data processing, curve fitting, and calculation of the fluorescence lifetime were all done using Symphotime 64 software (Picoquant). The longest lifetime with the higher fit amplitude τ1 was used to report membrane tension. A longer lifetime means more tension in the membrane (48).

### 2.13 Statistical analysis

All data presented were based on greater than three independent observations and the inclusion of biological replicates for *in vitro* analysis. The graphs represent the mean ± standard error. All statistical analyses were performed using Prism 10.1.2 (GraphPad). The difference between the two groups was assessed using paired or unpaired Student’s t-test when parametric statistical testing was applicable. When multiple groups were compared, a one-way analysis of variance followed by the Tukey post hoc test was applied. The p-value < 0.05 was considered a statistically significant difference between the test and control.

## 3. Results

### 3.1 MβCD and CHOL modulate cholesterol levels in HTM cells

Cyclodextrins (CDs) are cyclic oligosaccharides consisting of α-(1–4)-linked D-glucopyranose units, which contain a hydrophobic cavity and encapsulate various hydrophobic molecules. CDs typically exist as hexamers (αCDs), heptamers (βCDs) or octomers (γCDs). The degree of polymerization decides the size and affinity of the hydrophobic cavity (50). Among them, βCDs show the highest affinity for inclusion of cholesterol and the most efficient in extracting cholesterol from cell membranes (51). To improve the water solubility of βCDs, they are available in the methylated forms called MβCD (52), and acute cholesterol depletion by MβCD is the most commonly used method in many studies (40, 53). Due to the high affinity of MβCD for cholesterol, it can also be used to generate cholesterol inclusion complexes (CHOL) and donate cholesterol to the cell membrane (54). The experimental paradigm utilized in this study, is depicted in **Figure 1A**. “Cholesterol depletion” denotes treatment of HTM cells using 10 mM MβCD for 1h to remove membrane cholesterol. “Cholesterol enrichment” denotes treatment of HTM cells using 100 μM CHOL for 1 h to increase cellular cholesterol. “Cholesterol replenishment” denotes initial removal of cholesterol followed by supplementation. We confirmed that 10 mM MβCD and 100 μM CHOL treatment for 1 h removed and increased cholesterol respectively in HTM cells using Filipin staining (**Figure 1B**) and a free-cholesterol quantification kit (**Figure 1C**). Filipin is a naturally fluorescent polyene antibiotic that binds to free cholesterol but not esterified sterols (55). Filipin staining showed that under the control condition, cholesterol was distributed on the cell membrane, and cholesterol puncta were identified (denoted by white arrows) (**Figure 1B**). Compared to the control, 10 mM MβCD treatment for 1h decreased free cholesterol distribution, and no cholesterol puncta was observed (**Figure 1B**). However, 100 μM CHOL treatment for 1 h increased cholesterol distribution, and more cholesterol puncta was observed (denoted by white arrows) (**Figure 1B**). To further confirm the observations obtained using filipin staining, a free-cholesterol quantification kit was utilized and the result shows that compared to the control, 10 mM MβCD treatment for 1 h significantly decreased free cholesterol levels in HTM cells (n = 4, p = 0.03, **Figure 1C**). On the other hand, 100 μM CHOL treatment for 1 h significantly increased free cholesterol levels in HTM cells (n = 4, p = 0.0006, **Figure 1C**). These results indicate that using MβCD and CHOL can modify the cholesterol levels in HTM cells, laying the foundation for further experiments.

### 3.2 Actin cytoskeleton remodeling is dependent on cholesterol levels in HTM cells

Prior studies have highlighted the significance of cholesterol in the regulation of cell membrane properties, maintenance of membrane tension, actin cytoskeleton polymerization, and cell-ECM interactions (26, 56). However, the effects of cholesterol on the actin cytoskeleton are closely dependent on the cell types (57–60). To check the effects of cholesterol levels on the actin polymerization in HTM cells, changes in F-actin fiber formation were observed using SiR-Actin based live cell imaging. First, we observed changes in actin before and during 10 mM MβCD or 100 μM CHOL treatments for 1 h. Live cell imaging demonstrated that compared to pretreatment (0 min), the F-actin staining gradually decreased after MβCD treatment (**Figure 2A**). Quantitative image analysis using ImageJ measuring the F-actin fluorescent intensity in atleast 4 randomly chosen equal sized ROIs in each image (shown as boxes in the representative images, **Figure 2A**), confirmed that compared to the pretreatment (0 min), MβCD treatment for 20 min significantly decreased F-actin fluorescent intensity (n = 4, p = 0.005, **Figure 2B**), and continued through the 60 min that we observed. The fluorescent intensity of individual F-actin fibers was significantly decreased at 60 min compared to the pretreatment (0 min) (n = 57, p = 0.0001, **Figure 2C**). Therefore, confirming that cholesterol removal strongly induced actin depolymerization in HTM cells. Conversely, cholesterol enrichment exhibited gradual increase in F-actin staining every 10 min until the observed 60 min compared to pretreatment (0 min) (**Figure 2D**). Quantitative F-actin fluorescent intensity analysis verified the significant increase every 10 minutes (n = 4, p = 0.001, **Figure 2E**) compared to the pretreatment (0 min). Individual F-actin fiber intensity significantly increased after CHOL treatment for 60 min compared to pretreatment (0 min) (n = 48, p = 0.0001, **Figure 2F**). To better understand the reversibility of cholesterol mediated effects on the actin cytoskeleton, HTM cells were first treated with 10 mM MβCD for 60 min to deplete cholesterol, then washed thrice and replenished with 100 μM CHOL for another 60 min. The dynamic changes in actin cytoskeleton pretreatment and posttreatment were recorded using live cell imaging. As evident from the live cell imaging video (**Supplementary Video 1**), MβCD treatment gradually depolymerized actin as visualized by loss of F-actin fibers. Replenishing CHOL aided repolymerization of the actin fibers. The snapshots of the actin loss due to MβCD treatment for 60 min and the appearance of actin fibers due to CHOL replenishment are shown in **Figure 2G**. The quantitative changes in actin dynamics based on F-actin intensity compared between pretreatment (0 min), MβCD treatment and CHOL replenishment established that F-actin fluorescent intensity significantly decreased (n = 4, p < 0.05) through 60 minutes of cholesterol removal which was significantly reversed by cholesterol replenishment (n = 4, p < 0.05) (**Figure 2H**).

**Figure 2.**
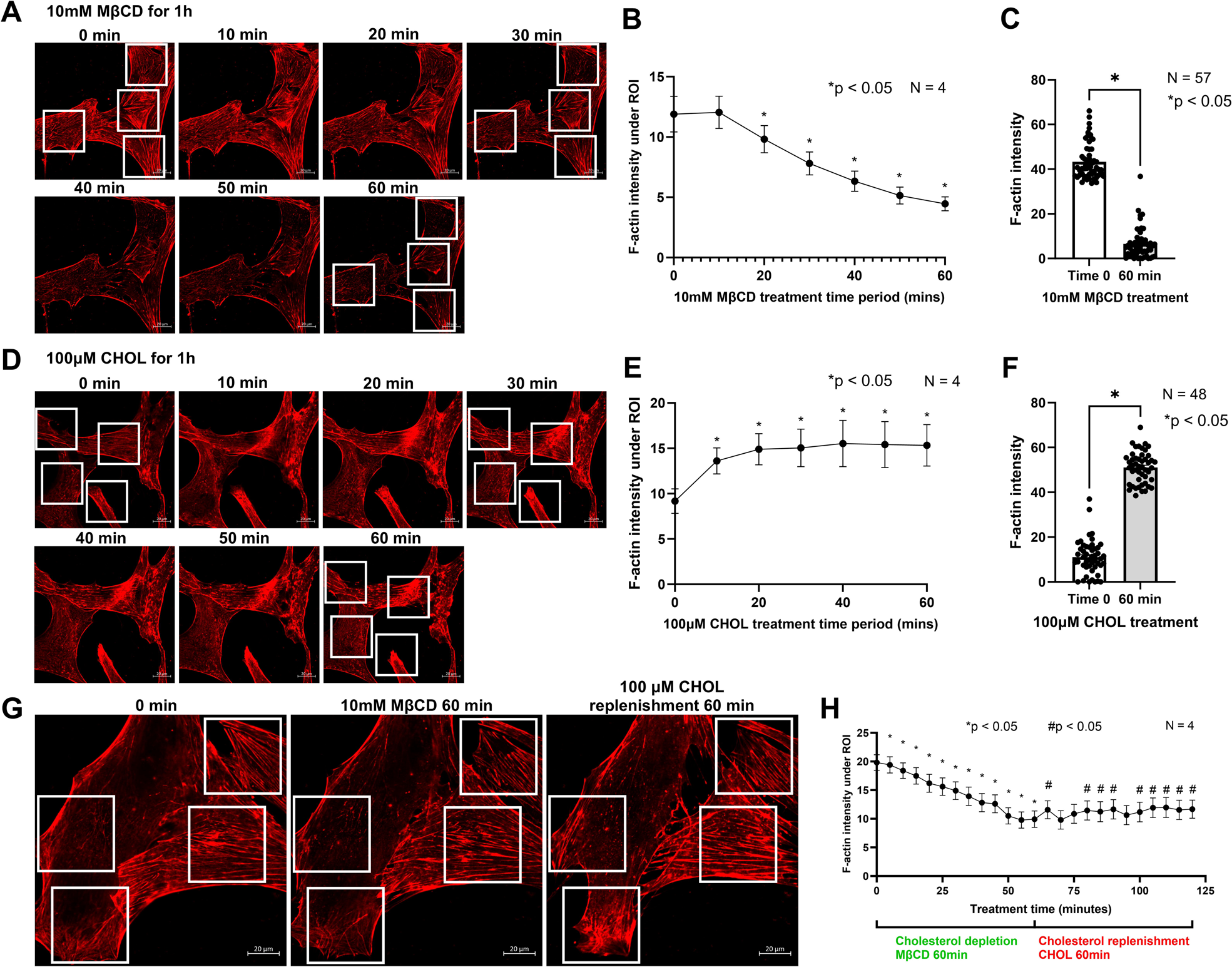
Cholesterol levels determine F-actin polymerization in HTM cells. (A) Live-cell imaging of HTM cells before and during 10 mM MβCD treatment for 60 min (captured every 10 min) to record dynamic changes of F-actin fibers stained with SiR-Actin. Compared to pretreatment (0 min), cholesterol depletion by MβCD gradually decreased F-actin fibers inside the cell. (B) F-actin fluorescent intensity in four ROIs (denoted in live-cell images) was analyzed, and compared to pretreatment (0 min), the F-actin intensity was significantly decreased after 20 min treatment and sustained until 60 min. (C) Individual F-actin fiber fluorescent intensity was compared between pretreatment (Time 0) and after MβCD treatment for 60 min. Compared to pretreatment, MβCD treatment for 60 min significantly decreased F-actin fibers fluorescent intensity. (D) Live-cell imaging of HTM cells before and during 100 μM CHOL treatment for 60 min (captured every 10 min) to record F-actin fibers dynamic changes. Compared to pretreatment (0 min), cholesterol enrichment by CHOL gradually increased F-actin fibers inside the cell. (E) F-actin fluorescent intensity in four ROIs (denoted in live-cell images) was analyzed, and compared to pretreatment (0 min), the F-actin intensity was significantly increased after 10 min treatment and sustained until 60 min. (F) Individual F-actin fiber fluorescent intensity was compared between pretreatment (Time 0) and after CHOL treatment for 60 min. Compared to pretreatment, CHOL treatment for 60 min significantly increased F-actin fibers fluorescent intensity. (G) Live-cell imaging of HTM cells before treatment (0 min), after cholesterol depletion by 10 mM MβCD treatment for 60 min, and after cholesterol replenishment by 100 μM CHOL treatment for another 60 min. Compared to pretreatment (0 min), MβCD treatment for 60 min decreased F-actin fibers in HTM cells, and cholesterol replenishment restored fibers. (H) F-actin fluorescent intensity in four ROIs (denoted in live-cell images) was analyzed, and compared to pretreatment (0 min), the F-actin intensity was significantly decreased after 5 min MβCD treatment and sustained until 60 min. Compared to 60 min after MβCD treatment, cholesterol replenishment by CHOL treatment (60 – 120 min) significantly increased F-actin intensity but cannot fully recover to the pretreatment level (0 min). Values represent the mean ± SEM, where n = 4-57. *p<0.05 was considered statistically significant when compared to pretreatment (0 min or Time 0). #p<0.05 was considered statistically significant when compared to the end time point of 60 min MβCD treatment.

Interestingly, the F-actin fluorescent intensity couldn’t fully recover it to the pretreatment level (0 min) (**Figure 2H**). These findings solidify the crucial role of cellular cholesterol levels in dictating F-actin polymerization and stability.

### 3.3 Cholesterol requires free actin monomers to induce F-actin formation

After confirming that cholesterol levels modulate actin polymerization in HTM cells, we determined the mechanism of cholesterol-induced F-actin dynamics. Firstly, we performed IF analysis post MβCD and CHOL treatments to study the F-actin (in green) polymerization and FA vinculin (in red) localization dynamics. Cholesterol removal using MβCD resulted in marked decrease in F-actin fibers and the distribution of vinculin at the focal edges whereas induced greater cytoplasmic accumulation (second row, **Figure 3A**) compared to the control condition (first row, **Figure 3A**). However, cholesterol enrichment showed observable increase in the actin polymerization and the thickness of F-actin fibers along with more vinculin distributed at the edges of F-actin fibers (denoted with black arrows, third row, **Figure 3A**). Thus, suggesting an increased association of actin with membrane via cell adhesive interactions. Compared to cholesterol depletion (MβCD), cholesterol replenishment (MβCD+CHOL) resulted in increased F-actin fibers and vinculin localization at the edges of F-actin fibers with less cytoplasmic accumulation (denoted with black arrows, fourth row, **Figure 3A**). Consistent with the removal of cholesterol using MβCD, inhibition of cholesterol biosynthesis using atorvastatin (100 μM) for 24 h on HTM cells, resulted in a marked decrease in F-actin fibers along with weakened distribution of vinculin at the actin edges (fifth row, **Figure 3A**).

**Figure 3.**
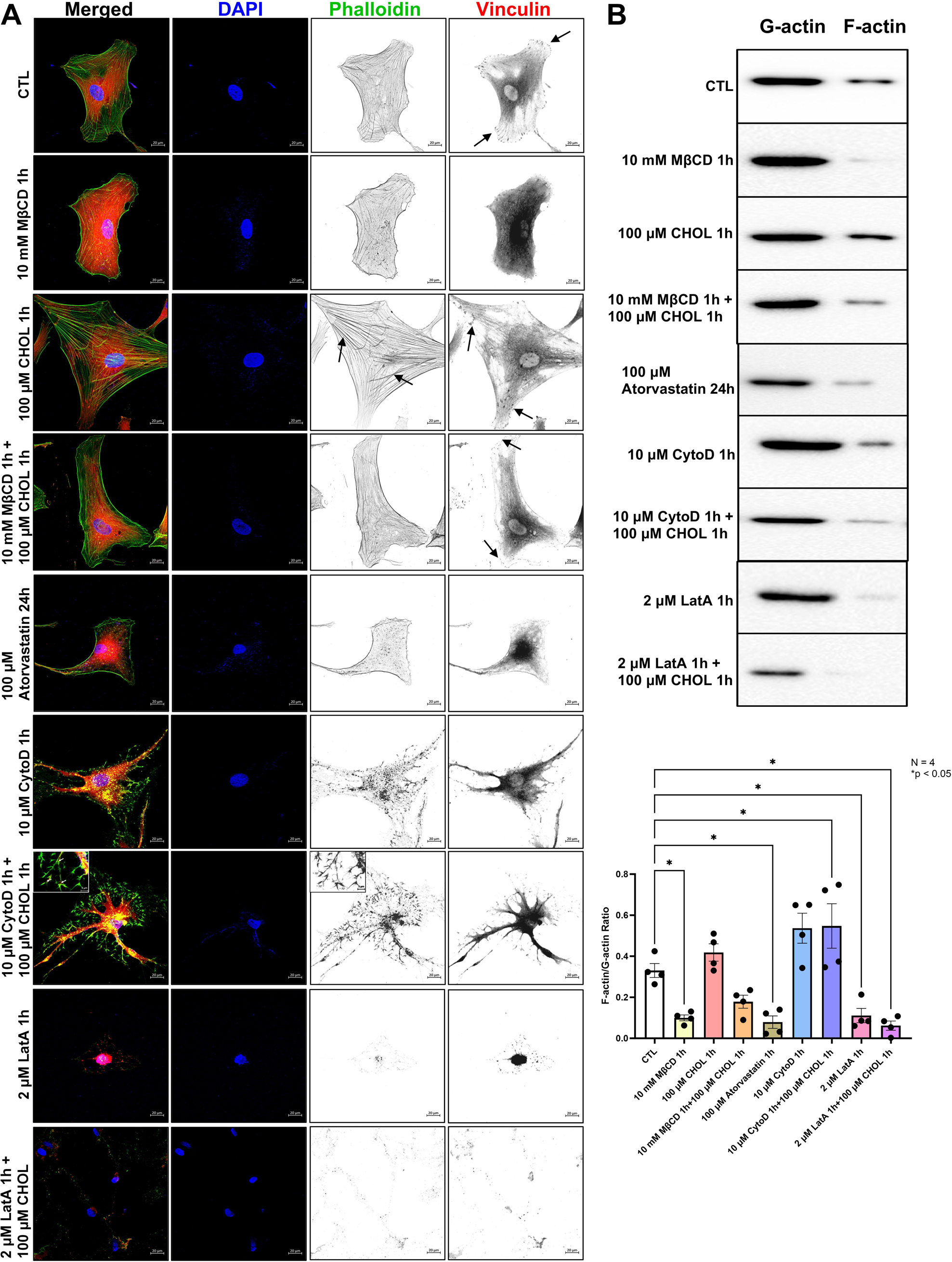
Cholesterol requires free G-actin monomers for actin polymerization in HTM cells. (A) IF imaging to explore the mechanism of cholesterol on actin polymerization by staining F-actin fibers with phalloidin and FA through checking vinculin distribution. Compared to the control condition (first row), cholesterol depletion by using MβCD decreased F-actin fibers with a sparse distribution of vinculin at the edges of F-actin fibers (second row). Compared to the control condition (first row), cholesterol enrichment by CHOL increased F-actin fibers and generated thicker fibers, along with more vinculin distributed at the edges of F-actin fibers (denoted by black arrow, third row). Compared to cholesterol depletion (second row), cholesterol replenishment (MβCD+CHOL) increased F-actin fibers and vinculin localization at the edges of F-actin fibers (denoted by black arrow, fourth row). Compared to the control condition (first row), cholesterol biosynthesis inhibitor atorvastatin caused a marked decrease in F-actin fibers with less vinculin distributed at the edges of F-actin fibers (fifth row). Compared to the control condition (first row), actin polymerization inhibitor CytoD decreased F-actin fibers with an F-actin remnants accumulation and cytoplasmic accumulation of vinculin in HTM cells (sixth row). CytoD combined with CHOL treatment (CytoD+CHOL) induced new F-actin fibers and actin branching formation with ∼70-degree angle (denoted in the inset, seventh-row third column), vinculin was distributed at the vertex of the F-actin branches (denoted by white arrows in the inset, seventh-row third column). Compared to the control condition (first row), LatA treatment decreased F-actin fibers and induced cytoplasmic accumulation of vinculin (eighth row). LatA combined with CHOL treatment (LatA+CHOL) didn’t restore F-actin polymerization in HTM cells (ninth row). (B) Quantitative analysis of F-actin/G-actin ratio after each treatment. Compared to the control condition, 1) MβCD treatment significantly decreased the ratio; 2) CHOL treatment increased the ratio but not significantly; 3) MβCD+CHOL increased the ratio slightly when compared to MβCD treatment, but still lower than the control condition; 4) atorvastatin decreased the ratio significantly; 5) CytoD increased ratio due to F-actin remnants accumulation; 6) CytoD+CHOL increased ratio significantly due to F-actin remnants accumulation and actin branching formation; 7) LatA caused a significant decrease in the ratio; and 8) LatA+CHOL didn’t resume the ratio and significantly lower than control condition. Values represent the mean ± SEM, where n = 4. *p<0.05 was considered statistically significant.

To decode the direct effects of cholesterol on actin polymerization in HTM cells, two actin polymerization inhibitors – cytochalasin D (CytoD) and latrunculin A (LatA) - were used. CytoD binds to the barbed ends of F-actin fibers with high affinity and inhibits both the polymerization and depolymerization of actin subunits at this end (61, 62). LatA directly binds to and inhibits the G-actin monomers binding to the F-actin barbed ends to disturb the actin polymerization (63). Compared to the controls, post 10 μM CytoD treatment for 1 h, we observed a marked decrease in F-actin fibers with the accumulation of F-actin remnants and cytoplasmic accumulation of vinculin (sixth row, **Figure 3A**). Strikingly, treatment of 100 μM CHOL for 1 h to HTM cells after a 1 h pretreatment of 10 μM CytoD induced new branches of F-actin fibers (Seventh row, **Figure 3A**). Interestingly, these newly formed F-actin fibers were actin-branches with an angle of ∼70-degree (denoted in the inset, seventh-row third column, **Figure 3A**). Of significance was also the recruitment of vinculin at the vertex of the F-actin branches (denoted by white arrows in the inset, seventh-row first column, **Figure 3A**). On the other hand, treatment with 2 μM LatA for 1 h caused an obvious decrease in F-actin fibers in HTM cells and induced cytoplasmic accumulation of vinculin (penultimate row, **Figure 3A**) compared to the control. Further, the provision of 100 μM CHOL in the presence of 2 μM LatA for 1 h did not restore F-actin polymerization and the focal adhesion recruitment (last row, **Figure 3A**). Thus, defining the significance of the availability of free G-actin monomers as the major determinant for cholesterol-induced F-actin polymerization and recruitment of focal adhesion.

Actin polymerization as a correlation to cholesterol availability was quantitatively assessed by measuring the F-actin/G-actin ratio. The immunoblotting and histogram in **Figure 3B** represents - 1) MβCD treatment significantly decreased the F-actin/G-actin ratio (n = 4, p = 0.02); 2) CHOL treatment increased the ratio, but not statistically significant (n = 4, p = 0.76); 3) cholesterol replenishment after MβCD treatment (MβCD+CHOL) caused a slight increase in the ratio when compared to MβCD treatment alone, but still lower than the control condition (n = 4, p = 0.23), which was consistent with results from live cell imaging and IF imaging that cholesterol replenishment partially rescue F-actin fibers; 4) atorvastatin treatment decreased the ratio significantly (n = 4, p = 0.01); 5) CytoD treatment and CytoD combined with CHOL treatment caused increased F-/G-actin ratio (n = 4, p = 0.05; n = 4, p = 0.04, respectively) which could be due to accumulation of F-actin remnants; 6) consistent with IF imaging results, both LatA treatment and LatA combined with CHOL treatment significantly decreased the ratio (n = 4, p = 0.03; n = 4, p = 0.007, respectively) (**Figure 3B**). All these results demonstrate that cholesterol levels are critical to maintain the actin polymerization status and recruitment of focal adhesions to the edges of the actin in HTM cells. The premise of such regulation relies on the unoccupied G-actin monomers.

Interestingly, actin-related protein 2/3 (Arp2/3) complex is involved in promoting branched actin filaments (64). To further understand how cholesterol regulates actin-branching after CytoD treatment, we checked the co-distribution of Arp2/3 complex and F-actin in HTM cells. Compared to the control condition, under CytoD treatment, F-actin fibers disappeared with accumulation of actin remnants (in magenta). Arp2/3 (in green) was disorganized, and vinculin (in red) was not distributed at the edge of F-actin fibers (**Figure 4A**). CytoD combined with CHOL treatment induced actin-branching (in magenta), and Arp2/3 (in green) and vinculin (in red) were localized at the vertices and actin edges of the actin branches respectively (**Figure 4A** and denoted by white arrows in zoomed-in images of ROIs, **Figure 4B**). Thus, defining the role of Arp2/3 in cholesterol-promoted actin-branching formation and vinculin maturation.

**Figure 4.**
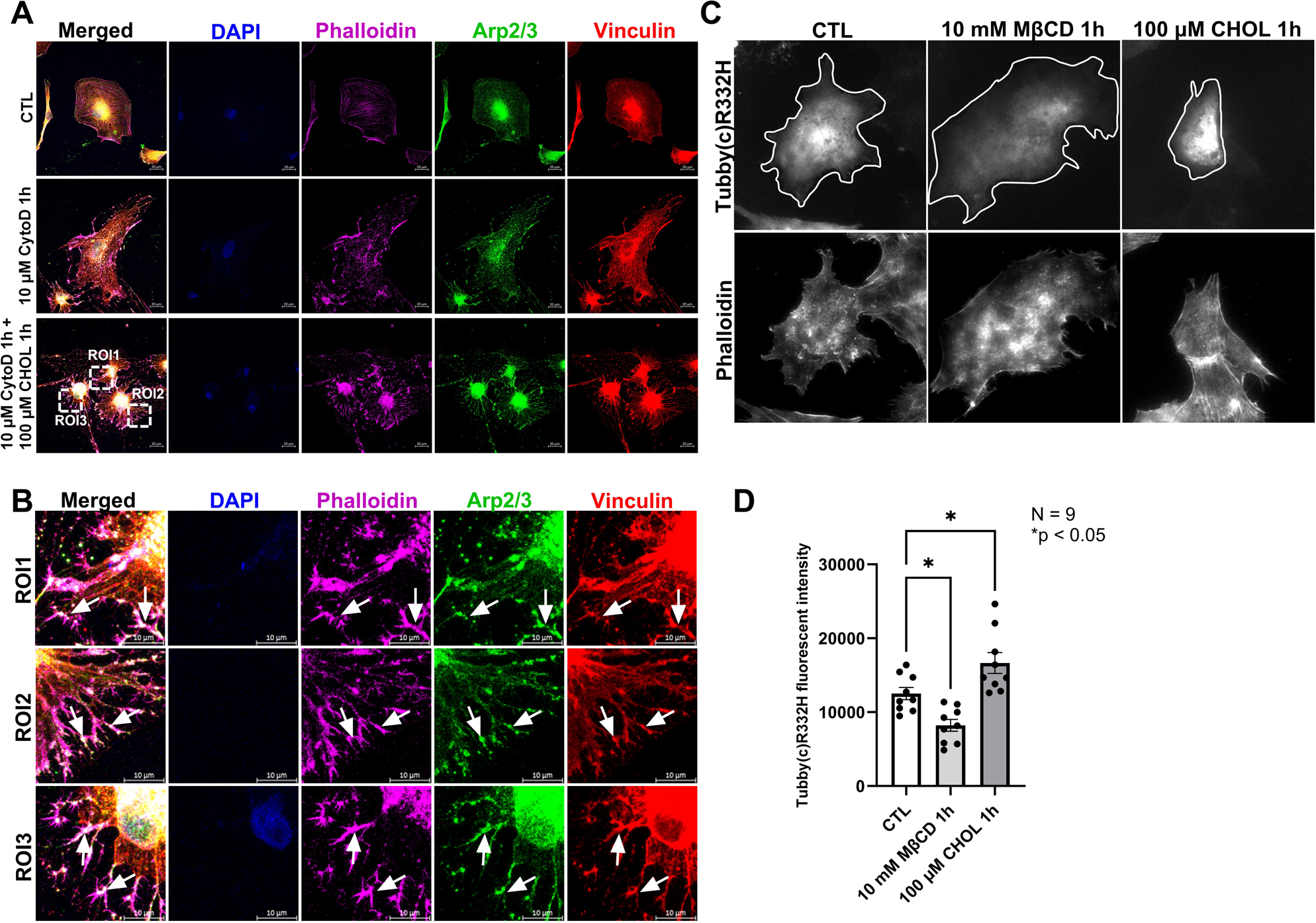
Cholesterol requires Arp2/3 and PIP2 to form actin fibers and branches. (A) and (B) IF to check cholesterol levels on actin-binding protein Arp2/3 distribution in HTM cells after CytoD and CytoD combined with CHOL treatments. (A) Compared to the control condition, CytoD treatment decreased F-actin fibers with an accumulation of actin remnants (in magenta), Arp2/3 (in green) was disorganized, and vinculin (in red) was not localized at the edges of F-actin fibers. CytoD+CHOL treatment induced F-actin polymerization and actin branching formation, with Arp2/3 and vinculin localized at the vertex of the actin branches. (B) Zoomed-in images of the chosen ROIs in the image of CytoD+CHOL treatment. CytoD+CHOL induced actin branching formation, and Arp2/3 and vinculin were located at the vertex at the actin branches (denoted by white arrows). (C) TIRF imaging of PIP2 dynamic changes on the cell membrane using Tubby(c)R332H plasmid transfection. Compared to the control condition, MβCD treatment decreased PIP2 levels on the cell membrane, and CHOL treatment increased the levels (cell membrane area was circled by white line). Phalloidin was used to identify cell shapes. (D) quantitative analysis of PIP2 changes on the cell membrane after treatments. Compared to the control condition, MβCD treatment significantly reduced Tubby(c)R332H fluorescent intensity on the cell membrane, and CHOL treatment significantly increased its intensity. Values represent the mean ± SEM, where n = 9. *p<0.05 was considered statistically significant.

### 3.4 Cholesterol modulates actin polymerization and cell adhesive interactions via phosphatidylinositol 4,5-bisphosphate (PIP2)

PIP2 is an essential lipid class present on cell membranes involved in the regulation of actin dynamics (65). Prior evidence suggests that membrane PIP2 regulates the function of many actin-binding proteins including formin, cofilin, and Arp2/3 (65, 66). PIP2 also binds to many FA proteins, such as vinculin, talin, and focal adhesion kinase (FAK), and serves as a linkage to FA and actin-binding proteins (65). Interestingly, previous studies found that cholesterol levels can regulate PIP2 on the cell membrane in a cell type-dependent manner (67–69). However, the effect of cholesterol levels on PIP2 in TM has not been explored. Due to the established connection between cholesterol levels and PIP2, as well as the significant function of PIP2 on actin dynamics and actin-binding proteins, we investigated the role of PIP2 on cholesterol-mediated actin changes in TM cells. NTM cells were transfected with pEGFP-N1-Tubby(c)R332H to indicate the changes of PIP2 on the cell membrane (44). TIRF imaging revealed cholesterol depletion using MβCD decreased PIP2 distribution on the NTM cell membrane (cell membrane area circled by white line, **Figure 4C**). In contrast, cholesterol enrichment increased PIP2 distribution on the NTM cell membrane (cell membrane area circled by white line, **Figure 4C**) compared to the control. Phalloidin staining was used to identify the cells. Quantitative analysis confirmed that cholesterol depletion (MβCD) significantly decreased the fluorescent intensity of Tubby(c)R332H on the cell membrane (n = 9, p = 0.01), and cholesterol enrichment (CHOL) significantly increased fluorescent intensity (n = 9, p = 0.02) (**Figure 4D**). This study demonstrates that cholesterol modulates PIP2 levels at the membrane, regulating dynamic actin polymerization and depolymerization.

### 3.5 Cellular cholesterol dictates the activation and distribution of integrins and matrix interactions

Cholesterol regulates cell-matrix interactions by modifying integrin sequestration, signaling, and adhesion (35). Cholesterol is critical in forming lipid rafts on the cell membrane, and the sequestration and recruitment of integrins in lipid rafts are required for integrins activation and signaling (70, 71). To understand the effect of cholesterol on cell-ECM interactions in HTM cells, we evaluated the qualitative distribution of total and active integrin α5β1 and αVβ3 using IF. We found that perturbation of cellular cholesterol - depletion (MβCD) and enrichment (CHOL) - had very little influence on the distribution of total integrin α5 and β1 (in red) in HTM cells (**Figure 5A and 5B**). On the other hand, activated integrin α5β1 (in red) was distributed at the edges of F-actin fibers (in green) in the baseline control condition (denoted by black arrow, **Figure 5C**). Cholesterol depletion (MβCD) decreased the overall distribution of activated α5β1 (**Figure 5C**). Whereas cholesterol enrichment (CHOL) modified the arrangement of activated α5β1 aligning more at the focal edges but pronounced changes were not seen compared to baseline (denoted by black arrow, **Figure 5C**). In the case of integrin αVβ3 and activated αVβ3 (in red) (**Figure 5D and 5E**) they decorated at the edges of F-actin fibers and near the cell membrane in the control condition (denoted by black arrow). Cholesterol depletion (MβCD) decreased their distribution at the edges of the cells with a cytoplasmic accumulation whereas cholesterol enrichment (CHOL) increased its distribution on the membrane as well as at the edges of F-actin fibers (denoted by black arrow, **Figure 5D and 5E**). Thus, providing a strong insight into the influence of cholesterol on the topographical distribution αVβ3 and α5β1 and their activation status in HTM cells.

**Figure 5.**
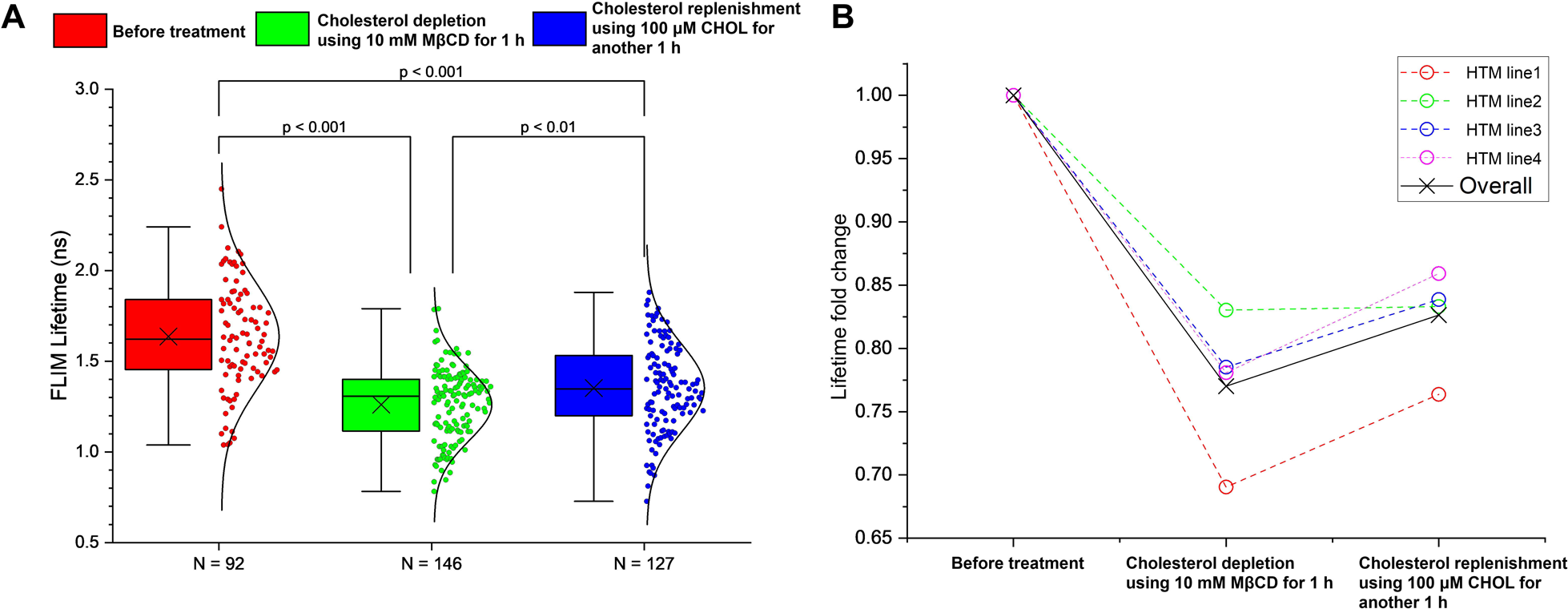
Cholesterol regulates localization of integrins and provides cellular force to sense substrate stiffness. (A) and (B) IF imaging of integrin α5 and β1 after cholesterol depletion and enrichment. Compared to the control condition, MβCD and CHOL treatments didn’t cause obvious changes in the distribution of integrin α5 and β1 in HTM cells. (C) IF imaging of activated integrin α5β1 distribution after cholesterol depletion and enrichment. Compared to the control condition, cholesterol depletion (MβCD) decreased the overall distribution of activated α5β1, whereas cholesterol enrichment (CHOL) modified the arrangement of activated α5β1 aligning more at the focal edges but pronounced changes were not seen compared to the control condition. (D) IF imaging of integrin αVβ3 after cholesterol depletion and enrichment. Compared to the control condition, MβCD treatment induced a marked decrease in the distribution of integrin αVβ3, and CHOL treatment caused increased distribution at the edges of F-actin fibers. (E) IF imaging of activated integrin αVβ3 after cholesterol depletion and enrichment. Compared to the control condition, MβCD treatment decreased the distribution of activated integrin αVβ3 at the edges of F-actin fibers with a cytoplasmic accumulation, and CHOL treatment increased its distribution at the edges of F-actin fibers. (F) TFM analysis of the traction force generated between HTM cells with COL1. The graph shows that compared to the control condition, MβCD treatment significantly reduced the traction forces, CHOL treatment didn’t cause a significant change, but it was significantly higher than MβCD treatment. Cholesterol replenishment (MβCD+CHOL) restored the forces, and it was significantly higher than cholesterol depletion (MβCD). (G) TFM analysis of the traction forces generated between HTM cells with COL4. The graph shows that compared to the control condition, MβCD treatment significantly reduced the traction forces, CHOL treatment didn’t cause a significant change in forces, but it was significantly higher than MβCD treatment. Cholesterol replenishment (MβCD+CHOL) restored the forces, and it was significantly higher than cholesterol depletion (MβCD), but it was still significantly lower than the control condition. (H) TFM analysis of the traction forces generated between HTM cells with FN. The graph shows that compared to the control condition, MβCD treatment significantly reduced the traction forces, CHOL treatment didn’t cause a significant change in forces, but it was significantly higher than MβCD treatment. Cholesterol replenishment (MβCD+CHOL) restored the forces, and it was significantly higher than cholesterol depletion (MβCD). (I) TFM analysis of the traction forces generated between HTM cells with laminin. The graph shows that compared to the control condition, MβCD treatment significantly reduced the traction forces, CHOL treatment didn’t cause a significant change in forces, but it was significantly higher than MβCD treatment. Cholesterol replenishment (MβCD+CHOL) restored the forces, and it was significantly higher than cholesterol depletion (MβCD). Values represent the mean ± SEM, where n = 6-10. *p<0.05 was considered statistically significant.

Integrin activation is critical to engage with specific ECM components like fibronectin, collagen, and laminin, and communicate with actin cytoskeleton. This communication generates forces applied to a surface by adherent cells. By measuring these forces in terms of traction force, we evaluated the effect of cholesterol on the strength of connections between HTM cells on different ECM substrates. Using printed microarrays/islands of collagen 1 (COL1), collagen 4 (COL4), fibronectin (FN), and laminin on 6 kPa polyacrylamide substrates, HTM cells were seeded onto these islands. TFM was utilized to measure the cellular forces exerted by calculating the displacement field before and after the relaxation of cellular forces (46). Overall, cholesterol depletion (MβCD) significantly decreased the traction force exerted by the cells on all ECM tested – COL1 (n = 10, p = 0.0002); COL4 (n = 9, p = 0.000001); FN (n = 6, p = 0.04); and laminin (n = 7, p = 0.002) (**Figure 5F, 5G, 5H, and 5I**). Though cholesterol enrichment (CHOL) didn’t cause significant changes in the forces compared to controls, it was significantly higher than cholesterol depletion for cells seeded on all the four ECMs – COL1 (n = 10, p = 0.0005); COL4 (n = 9, p = 0.00004); FN (n = 7, p = 0.004); and laminin (n = 7, p = 0.0006), cholesterol replenishment (MβCD+CHOL) significantly increased cell forces when comparing to cholesterol depletion (MβCD) – COL1 (n = 10, p = 0.001); COL4 (n = 9, p = 0.0007); FN (n = 9, p = 0.01); and laminin (n = 8, p = 0.007) (**Figure 5F, 5G, 5H, and 5I**). Based on these, we provide direct evidence that the cellular cholesterol content influences the traction force in a similar manner irrespective of the ECM substrate though the magnitude of influence varies. Thus, proving that TM cholesterol is a sine qua non in regulating cell-matrix interactions via integrins.

### 3.6 Changes in cellular cholesterol levels influence the HTM membrane tension and fluidity

As one of the abundant lipids in the cell membrane, cholesterol levels regulate cell membrane tension in various cell types (27, 72). We investigated the role of cholesterol in regulating cell membrane properties using a live cell fluorescent membrane tension probe Flipper-TR (48). Followed by the treatment of HTM cells with Flipper-TR, which can specifically insert into the cell membrane, cholesterol depletion was achieved by MβCD treatment followed by cholesterol replenishment using CHOL treatment. Using FLIM live cell imaging of HTM cells, the Flipper-TR fluorescence lifetimes in baseline/control condition (before treatment), after cholesterol depletion, and after cholesterol replenishment were acquired. In comparison to the baseline, cholesterol removal significantly decreased the fluorescence lifetime (n = 146, p = 0.000001) (**Figure 6A**). Not surprisingly, compared to cholesterol removal, replenishing cholesterol significantly increased the fluorescence lifetime (n = 127, p = 0.005), but was significantly lower than the baseline condition (n = 127, p = 0.000001) (**Figure 6A**). To derive the lifetime fold change in each HTM line utilized, we normalized the lifetime under different treatments to the baseline in each HTM cell line. Here, we observed that the lifetime average dropped by 30.95%, 16.96%, 21.48%, and 21.92% in each HTM cell line after cholesterol removal with an average drop by 22.98% (**Figure 6B**). Upon cholesterol replenishment, the lifetime average improved by 7.32%, 0.28%, 5.36%, and 7.83% in each HTM cell line averaging an improvement of 5.62% (**Figure 6B**). These findings indicate that cholesterol depletion significantly lowered HTM cell membrane tension and increased fluidity. Interestingly cholesterol replenishment partially restored membrane tension and decreased fluidity. Thus, signifying the key role of cholesterol in modifying HTM cell membrane properties.

**Figure 6.**
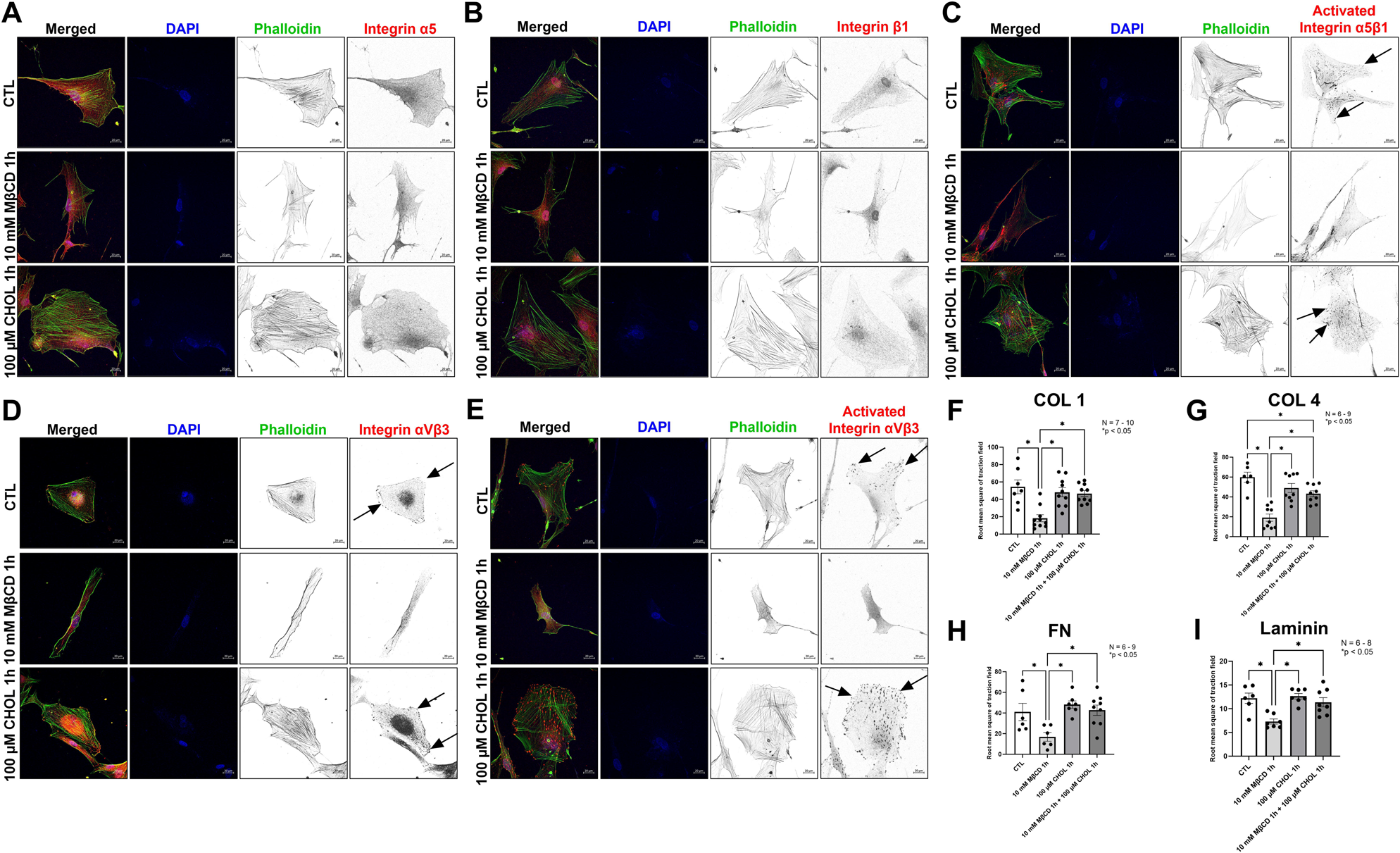
Cellular cholesterol levels contribute to the TM membrane tension and fluidity. FLIM analysis of Flipper-TR lifetime to indicate changes in HTM cells membrane tension and fluidity before (pretreatment), after cholesterol depletion (MβCD treatment), and after cholesterol replenishment (CHOL treatments). (A) FLIM analysis shows that compared to the baseline (pretreatment), cholesterol removal by MβCD significantly decreased Flipper-TR lifetime, indicating decreased cell membrane tension and increased fluidity; cholesterol replenishment by CHOL significantly increased lifetime compared to cholesterol removal, but still significantly lower than the control level. (B) Lifetime fold change in four HTM cell lines (biological replicates). The graph shows that compared to the baseline (pretreatment), cholesterol removal decreased lifetime by 30.95%, 16.96%, 21.48%, and 21.92% in each cell line, respectively. The average decrease was 22.98%. Compared to cholesterol removal, cholesterol replenishment increased the lifetime by 7.32%, 0.28%, 5.36%, and 7.83% in each cell line, respectively. The average increase was 5.62%. Values represent the mean ± SEM, where n = 92-146.

## 4. Discussion

Hyperlipidemia is associated with an increased risk of elevated IOP and glaucoma with poor mechanistic understanding (5, 6, 73, 74). Systemic cholesterol levels are closely related to glaucoma occurrence and IOP elevation (3–6). Interestingly, cholesterol-lowering statins lower IOP and reduce the incidence of glaucoma (8, 38). Our understanding of how TM cholesterol modulates the combined effects of cell stiffness, actin organization, matrix adhesion, and membrane tension is surprisingly limited. This study elucidates how cholesterol dynamically regulates the PIP2-actin interface, thereby fine-tuning actin organization and membrane interactions at the cellular periphery. Further, we provide the first detailed evidence on how cholesterol tunes the orchestra of actin cytoskeleton, cell adhesions, integrin, and matrix interactions in TM cells. Prior research into statins showed disruption of actin dynamics centered around a proposed mechanism where statins inactivate Rho GTPases by preventing their essential isoprenylation. Additionally, a genome-wide association study explored that gene ARHGEF12 was associated with IOP, and this protein is closely correlated with RhoA/RhoA kinase pathway (75), as well as binds to ABCA1, which is a known cholesterol efflux protein and influences the risk of developing POAG (76). By perturbing the cholesterol levels, we identify the direct role of cholesterol in dictating actin dynamics in TM and provide compelling evidence that cholesterol levels act as a critical molecular rheostat to determine cell stiffness. In general, it has been documented that TM tissue stiffness depends on actin cytoskeleton-based cell contractility and ECM remodeling (77). Whereas the knowledge on TM cell stiffness provided by the interactions of the actin cytoskeleton with the plasma membrane lipids and membrane receptors communicating outside the cells with ECM is largely unknown to date (78, 79). After comparing and contrasting the effects of statins versus the direct removal of cholesterol, our data clearly elaborates that the removal of cholesterol has a rapid effect of destabilizing the cytoskeletal machinery. Interestingly, replenishing cholesterol is not able to completely reverse the actin to baseline suggesting that cholesterol removal erases the cellular “morphological” memory. We propose that cells take a new shape exhibiting a lower contractile force as cholesterol removal leaves a stronger effect on actin makeup and cell adhesive interaction including focal adhesions and integrin localization and activation in the cells. This makes cholesterol removal use as an interesting target to lower IOP. The requirement for free G-actin monomers for cholesterol-mediated actin polymerization highlights a unique regulatory mechanism. Moreover, the observation of both linear and branched actin networks in the presence of cholesterol and G-actin suggests diverse effects on actin filament assembly. Arp2/3 and focal adhesions were induced at the actin network’s vertices and leading edges, respectively. Thus, suggesting that cholesterol in concert with actin can aid in increasing filopodia and lamellipodia formation. In fact, this phenomenon was observed upon activation of SREBPs, which is known to increase cellular lipids (7). Interestingly though, recently, it was reported that inhibition of Arp2/3 decreased the aqueous humor flow rate (80). Therefore, it is important to evaluate the effect of cholesterol removal on the magnitude of aqueous humor outflow and IOP regulation.

Curiously, a previous study using MβCD to deplete cholesterol in HTM cells demonstrated that MβCD increased actin polymerization after treatment by facilitating agonist induced TRPV4 activation (40). This data surprised us as we found the opposite result in HTM cells. Employing a variety of experimental paradigms, our study consistently demonstrated that cholesterol depletion reduces actin-based cytoskeletal tension. Our results align with prior research showing statins relax HTM cells and limit actin polymerization, further supporting the link between cholesterol biogenesis and actin tension (37, 38). Cholesterol molecules wedge themselves between phospholipid molecules in the membrane, making it denser and less fluid (81). This increases the membrane’s rigidity and resistance to bending. Membrane lipids with long tails and saturated fatty acids can increase cell stiffness and decrease cell membrane fluidity whereas unsaturated lipids do the opposite since acyl-chain kinks prevent tight packing. Cholesterol interacts with sphingolipids, another type of lipid in the membrane, to form microdomains called “lipid rafts”. These rafts are even more rigid than the surrounding membrane, further contributing to increased stiffness. By understanding the role of PIP2 in cholesterol-mediated actin changes in TM, our research sheds light on the precise mechanisms by which cholesterol influences critical cellular processes like actin dynamics, adhesion, and matrix interactions. Typically, PIP2 is concentrated in cholesterol-rich microdomains on the cell membrane, such as caveolae and lipid rafts and changes in cholesterol levels on the cell membrane can generate dramatic effects on PIP2 levels and distribution (68, 69). PIP2 is also related to integrin activation by binding to multiple FA proteins and linking the FA and integrin signaling pathways (65). An optimal balance in ECM remodeling in JCT-TM region is needed for the generation of AH resistance, but excess ECM deposition results in increased resistance and elevated IOP (16, 17). In POAG, characteristic changes occur in the tissue structure of the TM AH outflow pathway (15) including excessive alterations in ECM organization in the JCT-TM region and accumulation of sheath-like plaque materials leading to altered stiffness (16, 77). Better understanding of lipid rafts in PIP2 signaling, cytoskeletal and ECM changes regulated by PIP2 in TM and their contribution for TM stiffness in TM physiology and pathobiology warrants a detailed study.

While we recently showed that inhibiting lipogenesis in TM reduced ECM genes, changes in ECM protein distribution, and fibril formation (7). The present study provides a direct link on how cholesterol plays a central cog in regulating the actin-focal adhesion-integrin link via PIP2. Integrins are known to be involved in TM biomechanics and glaucoma pathogenesis (20, 21). Integrins act as mechanotransducers in TM cells to respond to the extracellular environment (22), control matrix assembly by tethering fibronectin, and its activation also regulates actin cytoskeleton remodeling (20). Moreover, deletion of integrin β3 results in a significant reduction in IOP in animal models, and its activation increases the IOP (82). Our findings build a connection between cholesterol levels and integrin activation, further signifying the critical role of cholesterol in cell-ECM interactions in TM cells. Interestingly, TFM data shows that the response to cholesterol removal and supplementation of the four ECMs tested are very similar in magnitude. It is likely that when the cells adhere to the ECM and pull on these proteins, due to the thin 2D nature of the ECM, the cells pull on both the ECM and the acrylamide, and the acrylamide stiffness has a major impact on the cell response. Future studies are needed to understand how different ECM substrates affect the integrin expression and the traction force experienced by the cells.

Cholesterol can change the lipid organization of the plasma membrane. The presence of cholesterol in the membrane helps the phospholipid tails to align and pack closer together, generating membrane order increasing its tension and rigidity and lowering permeability (78). Increased TM tissue mechanical tension and stiffness are important for IOP elevation (16, 83). As one of the critical lipid components of the cell plasma membrane (27), we observed that cholesterol depletion significantly reduced cell membrane tension, and cholesterol replenishment partially restored the tension. This is in good agreement with the known effects of cholesterol on membrane tension and fluidity (56).

In conclusion, our study unveils cholesterol as a master regulator of TM cell mechanics, controlling actin dynamics, membrane interactions, and tension. The intricate link between cholesterol and PIP2 sheds light on how it tunes TM stiffness, connecting to clinical observations and suggesting that modulating cholesterol levels in TM cells offers a promising avenue for treating glaucoma.

## 5. Data availability statement

All data are contained within the manuscript. Any additional data will be provided upon request.

## 6. Acknowledgements

We acknowledge Dr. Nuria Morral, Dr. Benjamin Perrin, Dr. Gary Landreth, and Dr. Tim Corson from IUSM for their helpful discussions on the project. This project was supported by the National Institutes of Health/ National Eye Institute grant R01EY029320 (PPP), NIH/NIGMS grant R35GM147412 (JL), and NIH R01DK115747 (GHU). Award from the Ralph W. and Grace M. Showalter Research Trust and the Indiana University School of Medicine (PPP), Research Support Funds Grant (RSFG), Cohen AMD Research Pilot Grant (PPP), RPB Departmental Pilot Grant (PPP), Glick Research Endowment Funds (PPP), and Challenge grant from Research to Prevent Blindness to IU.

## 7. Conflict of Interest

The authors declare that they have no conflict of interest. The funders had no role in the design of the study; in the collection, analyses, or interpretation of data; in the writing of the manuscript, or in the decision to publish the results.

## 8. Author contributions

Conceptualization: PPP. Methodology - Cell cultures, immunofluorescence, immunoblotting: TW. TIRF and TFM imaging: HRCK, HR, TW. FLIM imaging: CP, TW. Formal analysis: TW, HRCK, CP, JL, GHU, and PPP. Investigation: TW, HRCK, CP, and PPP. Write-up and data curation: TW, HRCK, CP, JL, GHU, and PPP. Writing—original draft preparation: TW and PPP. Figure preparation: TW, HRCK, CP, and PPP. Visualization: TW and PPP: Supervision: PPP: Project administration: PPP: Funding acquisition: PPP, JL, GHU. All authors have read and approved the manuscript.

## 9. Ethics declarations

### Ethics approval

Ethical review and approval were waived for the use of cadaveric human eyes for the isolation of the TM cells from human cadaveric corneal rims for this study.

---

Supplementary video 1 legend:

Live-cell imaging of HTM cells captured every 5-minutes before and during cholesterol depletion using 10 mM MβCD for 1 h, and during cholesterol replenishment using 100 μM CHOL for another 1 h. F-actin fibers were stained with SiR-Actin. The video shows that MβCD treatment caused a gradual decrease in F-actin fibers in HTM cells (denoted by white arrow). After 1 h MβCD treatment, the cells were replenished with cholesterol by CHOL treatment, the video shows that cholesterol replenishment restored F-actin fibers in the HTM cells (denoted by white arrow). The time, SiR-Actin, and treatments (MβCD and CHOL) are labeled at the top left corner of the video.

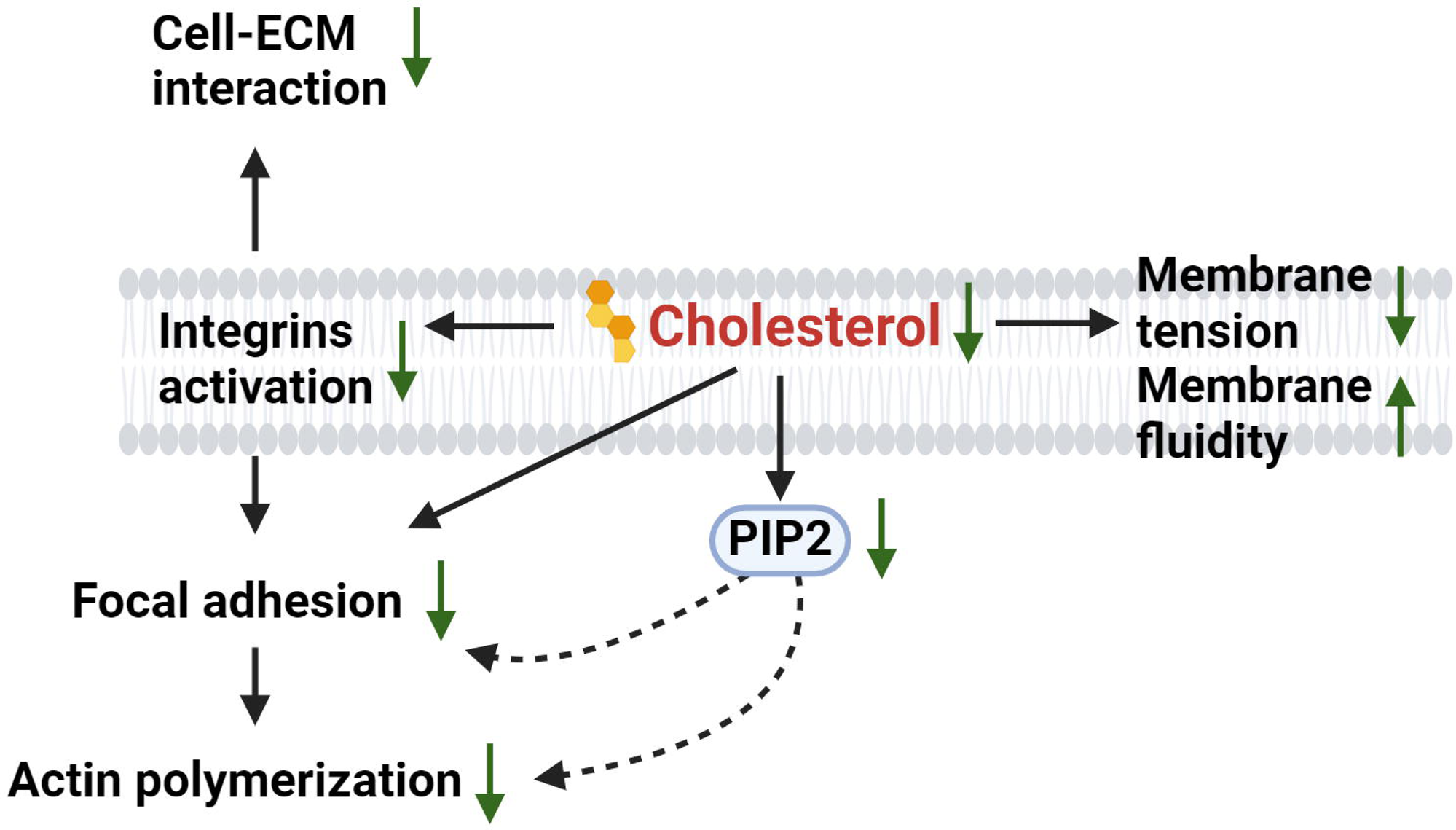

## Notes

### Competing Interest Statement

The authors have declared no competing interest.

